# Prioritizing candidate peptides for cancer vaccines by PEPPRMINT: a statistical model to predict peptide presentation by HLA-I proteins

**DOI:** 10.1101/2021.09.24.461596

**Authors:** Laura Y. Zhou, Fei Zou, Wei Sun

**Affiliations:** Department of Biostatistics, University of North Carolina at Chapel Hill; Public Health Science Division, Fred Hutchinson Cancer Research Center; Department of Biostatistics, University of Washington

**Keywords:** peptide-HLA association, Neural Network, mixture model, neoantigens, melanoma, cancer vaccine

## Abstract

Recent development of cancer immunotherapy has opened unprecedented avenues to eliminate tumor cells using the human immune system. Cancer vaccines composed of neoantigens, or peptides unique to tumor cells due to somatic mutations, have emerged as a promising approach to activate or strengthen the immune response against cancer. A key step to identifying neoantigens is computationally predicting which somatically mutated peptides are presented on the cell surface by a human leukocyte antigen (HLA). Computational prediction relies on large amounts of high-quality training data, such as mass spectrometry data of peptides presented by one of several HLAs in living cells. We developed a complete pipeline to prioritize neoantigens for cancer vaccines. A key step of our pipeline is PEPPRMINT (PEPtide PResentation using a MIxture model and Neural neTwork), a model designed to exploit mass spectrometry data to predict peptide presentation by HLAs. We applied our pipeline to DNA sequencing data of 60 melanoma patients and identified a group of neoantigens that were more immunogenic in tumor cells than in normal cells. Additionally, the neoantigen burden estimated by PEPPRMINT was significantly associated with activity of the immune system, suggesting these neoantigens could induce an immune response.

## 1 Introduction

Cancer immunotherapy harnesses the power of the human immune system to eliminate tumor cells and has become a new pillar of cancer treatment, in addition to chemotherapy, surgery, and radiation [1]. The successes of the existing immunotherapy solutions have motivated investigators to find alternative approaches to activate the immune system against cancer. One direction is developing cancer vaccines made of neoantigens, i.e., peptides harboring tumor-specific somatic mutations [2–4]. A cancer patient may have hundreds or thousands of somatic mutations and thus a crucial step for cancer vaccine production is prioritizing neoantigens that can induce an immune response. Once such neoantigens are identified, vaccines are manufactured using the peptides around the corresponding somatic mutations [3, 4]. Each vaccine may contain up to 20 such peptides and can be used alone or combined with other treatments, such as the popular immunotherapy check-point inhibitor treatment [5].

In most living cells, proteins are processed to peptides of certain lengths and transported to the cell surface to be presented by the human leukocyte antigen (HLA) proteins. Many computational methods have been developed to predict peptide presentation using the data generated by *in vitro* binding affinity assays [6–9]. These assays are low-throughput and do not reflect the restrictions due to peptide processing and transportation [9–14]. In contrast, the *in vivo* mass spectrometry (MS) method is high-throughput and recovers eluted peptides that reflect the restrictions of peptide processing and transportation [10]. Most of the MS data collected are from living cells that express up to six HLA-I proteins from the maternal/paternal copies of three highly polymorphic genes: HLA-A, HLA-B, and HLA-C [15]. Therefore, a major challenge of using MS data is identifying which corresponding HLA allele is associated to each eluted peptide. Although it is possible to produce single HLA allele MS data [12, 16], which we refer to as **SA data**, multi-HLA-allele MS data (or **MA data**) are much easier to generate and more abundant. Currently, methods that directly model MA data are lacking, and as the MA data source grows larger, these types of methods are critically needed.

Both SA and MA data share the challenge that training datasets may only cover a limited number of HLA alleles out of the thousands of discovered HLA alleles in the human population [17]. To address this challenge, information across HLA alleles are borrowed using a pan-specific approach where each HLA allele is represented by its sequence. Thus, the trained model can be applied for any HLA allele with sequence information [16, 18, 19]. To address the challenge of the ambiguity of peptide-HLA associations in MA data, some previous methods cluster the peptides un-supervisedly and assign each cluster to an HLA allele based on prior information [20–22]. The application of these methods are limited since prior information is only available for a small number of HLA alleles. Alternatively, NNAlign MA [19] takes a semi-supervised approach. NNAlign MA essentially uses a hard k-means clustering method, where each peptide is assigned to one and only one cluster (or HLA) using a pan-specific neural network with binding affinity and SA data. Then, it iteratively updates the neural network and inferred pairings of the peptides and HLA alleles. NetMHCpan-4.1 is created by training NNAlign MA [19] on a more recent MA training data set.

Motivated by NetMHCpan-4.1, we propose a pan-specific mixture model named PEPPRMINT for multi-HLA-I allele mass spectrum data (i.e., MA data). Each mixture component of our model corresponds to an HLA allele and the density function of each mixture component is specified by a neural network. PEPPRMINT has several practical advantages against NetMHCpan-4.1. First, it has an explicit objective function, the likelihood of the mixture model, which helps monitor algorithm convergence. Second, PEPPRMINT accounts for the uncertainty of associating each peptide with an HLA allele by assigning the peptide to different HLA alleles with appropriate weights. In contrast, NetMHCpan-4.1 assigns each peptide to one and only one HLA allele. Third, as a mixture model, PEPPRMINT estimates mixture proportions and uses them to improve the accuracy of assigning a peptide to an HLA allele. This is important to incorporate because different HLA alleles may present very different numbers of peptides [20, 21], for example, due to the low expression of an HLA allele [20] or somatic mutations of an HLA allele [23]. In contrast, NetMHCpan-4.1 assumes the mixture proportions are the same for all the HLA alleles.

We have developed a computational pipeline to prioritize neoantigens for cancer vaccines using whole exome sequencing data. Starting with the raw data of fastq files, our pipeline maps sequence reads, calls and annotates somatic mutations, imputes HLA alleles, extracts peptide sequences around somatic mutations, and predicts their presentation by any HLA allele of a subject using PEPPRMINT or NetMHCpan-4.1. In a case study, we applied our pipeline to analyze the exome-seq data of 60 melanoma patients [24]. We demonstrated that for many somatic mutations, the mutated peptides and the corresponding wild-type peptides had similar chances to be presented by HLA proteins. We prioritized those somatic mutations of which the mutant copies were highly likely to be presented on cell surface while the wild type copies were much less likely to be presented, since these peptides should induce a strong and tumor-specific immune response. We also demonstrated that neoantigen burden derived from PEPPRMINT had a significant association with immune cytolytic activity, suggesting that the neoantigens prioritized by PEPPRMINT would induce an immune response.

## 2 Method

### 2.1 An overview of our pipeline and PEPPRMINT

We designed a pipeline to prioritize candidate peptides for cancer vaccines using whole exome-seq data from tumor samples and paired normal samples (Figure 1 (A)). Starting with fastq files, we mapped DNA sequences to the human reference genome (build hg38) by BWA [25]. After applying a series of pre-processing steps, such as tagging duplicated reads and calibrating quality scores [26], we called somatic mutations using both Mutect [27] and Strelka [28]. The intersection of these somatic mutation calls were annotated by ANNOVAR [29] to obtain the functional consequence of the mutations and their associated genes and proteins. We selected non-silent mutations (i.e., those that alter the proteome) and extracted peptide sequences around those mutations from both the tumor proteome and the normal proteome. Then, we applied our method, PEPPRMINT, and NetMHCpan-4.1 to score the associations between these peptides and any HLA of each subject, where each HLA was imputed by OptiType using exome-seq data [30].

**Figure 1:**
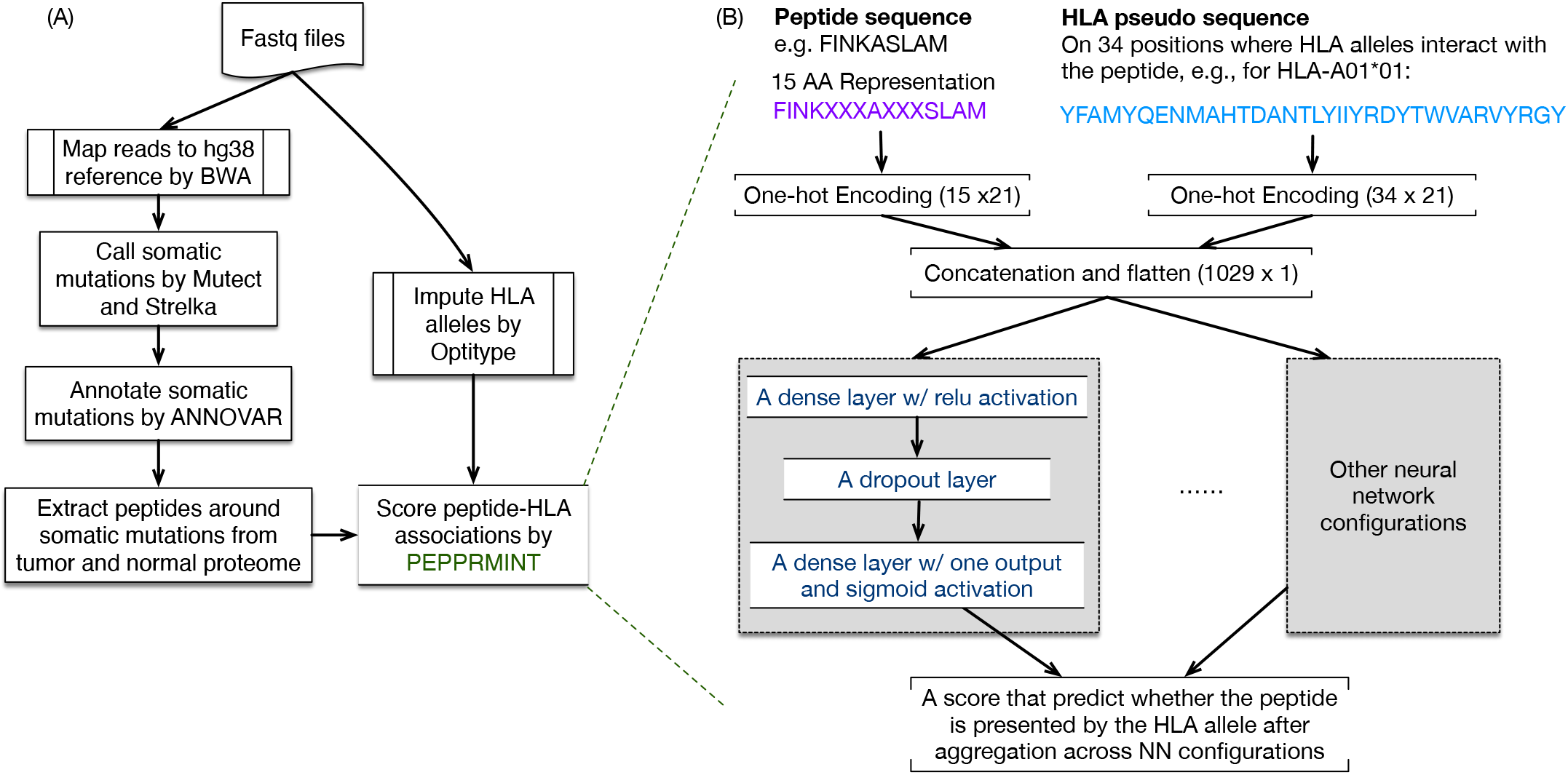
(A) An outline of our pipeline that prioritizes candidate peptides for cancer vaccine. (B) The neural network estimated by our PEPPRMINT method. Several versions of the neural networks (e.g., trained using different network configurations or input data) are aggregated together by taking the average of the neural network outputs.

While most of the components in our pipeline are based on existing methods, PEPPRMINT is new. PEPPRMINT estimates a neural network that predicts how likely a peptide is presented by an HLA allele using the amino acid (AA) sequence of the peptide and the pseudo-sequence of the HLA allele, which covers 34 AAs that have close contact with the presented peptides [31] (Figure 1 (B)). In order to train this model using MA data, we embedded this neural network within a mixture model framework, where each mixture component corresponded to an HLA allele and the neural network modeled the density function of each mixture component. Several components of this framework required careful investigation. The lengths of peptides vary from 8 to 15 AA, however a neural network requires the data input to have the same dimension. Therefore, peptides need to be transformed to a uniform length. Additionally, the configurations of the neural networks, such as the number of layers, the number of nodes, as well as the parameters for model training, need to be selected. We elaborate on these issues in the following sections, after providing more details of the input data and introducing our statistical framework.

When applying our method for cancer vaccine design, we prioritized personalized neoantigens for each cancer patient based on her/his somatic mutations and HLA alleles. We applied the trained neural networks to predict the associations between each HLA allele and each mutated/wild-type peptide, and took the maximum across the HLA alleles and the peptides covering the same somatic mutation. We used these predictions to rank neoantigens as candidates for a cancer vaccine.

### 2.2 Data

#### 2.2.1 Training Data

We trained our model using the same training data as NetMHCpan-4.1 [32] compiled from multiple sources and included both single-HLA-allele and multi-HLA-allele mass spectrum data (SA and MA data, respectively). There were around 200,000 and 350,000 positive peptides (binders, 8 to 15 AA long) for SA and MA data, respectively (Figure 2 (A)). Negative peptides (non-binders) were generated by randomly sampling the human proteome [12,32]. An equal number of non-binders were generated for each peptide length (5 times of the number of 9 AA binders). The number of peptides per HLA in the SA data had a wide range from 6 to more than 200,000, with median being around 4,000 (Figure 2 (B)), for 130 unique human HLAs. The MA data comprised of 105 unique samples, with three to six HLA alleles for each sample (Figure 2 (D)) and around 1,000 to 400,000 peptides per sample (Figure 2 (E)). Additional details of the training data are summarized in the Supplementary Materials.

**Figure 2:**
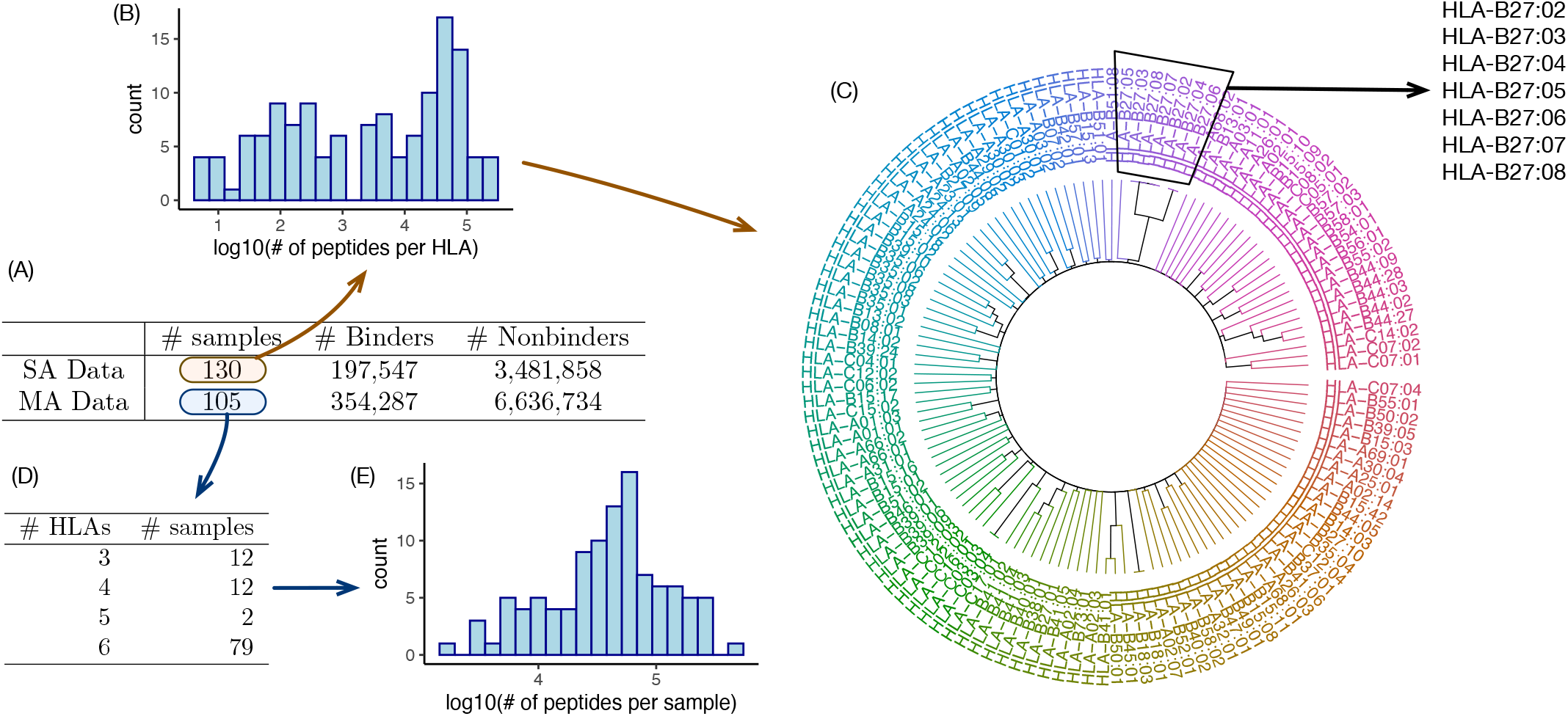
(A) Summary of single-HLA-allele mass spectormetery (SA) and multi-HLA-allele mass spectrometery (MA) training data. (B) The distribution of the number of peptides (including binders and non-binders) per HLA (or equivalently, per SA sample). (C) Hierarchical clustering of HLA alleles in SA data, where two HLA alleles have smaller distance if they shared more peptides than expected by chance. (D) The distribution of the number of HLAs per sample for MA data. (E) The distribution of the number of peptides per sample for MA data.

In the SA data, some binders were shared across HLAs. As we showed in the following analyses, such sharing reflected the biological reality that some HLA alleles are very similar. We quantified whether two HLA alleles shared more peptides than expected by chance through a hyper-geometric test, where a smaller p-value indicated that the two HLAs shared more binders than expected by chance. After truncating the p-value at a lower-end of 10^*−*10^, we further transformed the p-values to a distance measurement 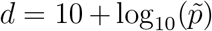, where 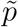 was the truncated p-value. We clustered the HLAs using this distance matrix and found those HLAs with smaller distances (i.e. more overlapping peptides) were similar in their DNA sequences, as reflected by their names (Figure 2 (C)). Such sharing of binders across HLAs showed that it might not be feasible to assign a peptide to one and only one HLA allele without ambiguity.

Each record in the SA data included three entries: peptide sequence, the corresponding HLA-I allele, and the binder status (i.e. binder = 1, non-binder = 0). Each record in the MA data included peptide sequence, the corresponding sample, and the binder status. Additionally, a separate file listed the specific HLAs for each sample. Both the SA and MA data were partitioned into 5 splits. We trained our model using each split separately and then aggregated the results by taking average.

#### 2.2.2 Testing Data

We evaluated PEPPRMINT and other methods using two testing datasets. The first test set was the MULTIALLELIC-RECENT data used by MHCflurry-2.0 [16]. This MA data was comprised of 20 experiments (i.e., 20 unique samples) from two studies. Each sample had 6 HLAs and the peptide lengths varied from 8 to 11 AA. Synthetic non-binders were randomly sampled peptides from the human proteome [16]. The total number of non-binders generated was ten times the number of binders, generated evenly across lengths 8-11. Peptides that were duplicated in the same sample were removed, which resulted in a total of 2,678,188 peptides (25,309 binders).

Though the MA test set better reflected the multi-allelic nature of cancer vaccine application than a SA test set, a second testing dataset comprised of SA peptides obtained from Sarkisova et al [33] and provided by NetMHCpan-4.1 [32] was used. After removing duplicated peptides associated with the same HLA, there were 946,008 peptides (45,416 binders) for 36 HLAs, varying from 8 to 14 AAs in length. Non-binders were added by extracting 8-14 AA peptides from source proteins of the binders.

### 2.3 Mixture model

Assume that binding peptides are collected for *n* samples. For the *i*-th sample with *K*_*i*_ HLA alleles and *J*_*i*_ peptides, let *x*_*ij*_ and *θ*_*ik*_ be the sequences for the *j*-th peptide and the pseudo sequence of 34 amino acids [31] for the *k*-th HLA allele, respectively, where *j* = 1, ..., *J*_*i*_ and *k* = 1, ..., *K*_*i*_. The total number of binders per sample, *J*_*i*_, usually varies from hundreds to tens of thousands, and *K*_*i*_, the number of unique HLA-I alleles, varies from 1 to 6. Since it is unknown which HLA allele presents the *j*-th peptide, we consider all possible *K*_*i*_ HLA alleles and model the likelihood of peptide presentation by a mixture model. The likelihood function for binders only is

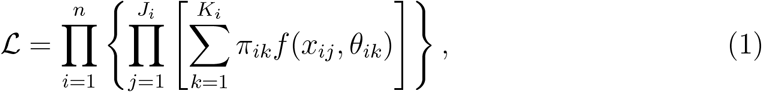

where ∑_*k*_ *π*_*ik*_ = 1 and *π*_*ik*_ is the probability that a randomly selected peptide is presented by the *k*-th HLA allele of sample *i*. Here, the function *f*(*x*_*ij*_, *θ*_*ik*_) is the probability that *x*_*ij*_ is presented by the *k*-th HLA allele (with pseudo sequence *θ*_*ik*_) and is modeled by a neural network. The output layer of this neural network uses a sigmoid activation function, *ϕ*(*z*) = 1*/*(1 + exp(−*z*)), which guarantees *f*(*x*_*ij*_, *θ*_*ik*_) falls between 0 and 1.

In order to train the neural network to distinguish not only the binders of different HLA alleles, but also binders versus non-binders, we introduce *L*_*i*_ non-binders into the model. Let *z*_*il*_ be the sequence of the *l*-th non-binder, which is randomly assigned to an HLA allele *k*_*l*_ of the set of all possible *K*_*i*_ HLAs for sample *i*. Then, the final likelihood function becomes

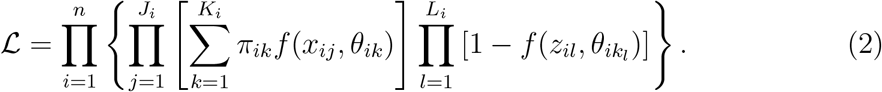

We estimate the parameters of this likelihood function by a standard EM algorithm. We first obtain an initial estimate of *f*(*, *θ*_*ik*_) using the single-allele MS data and assign the initial value of *π*_*ik*_ to be 1/*K*_*i*_. Then, the EM algorithm proceeds as follows. In the E-step, we evaluate the possibility that *x*_*ij*_ is presented by any one of the possible HLA alleles. For sample *i*, the probability the *j*-th peptide is presented by the *k*-th HLA allele is 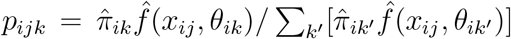. In the M-step, we estimate two sets of parameters: *π*_*ik*_ and *f*(*, *θ*_*ik*_) using the weighted input data for the neural network optimization. When estimating *π*_*ik*_, we can ignore the likelihood of non-binders and by standard EM algorithm results: 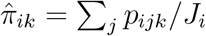. When estimating *f*(*, *θ*_*ik*_), the corresponding *Q* function is

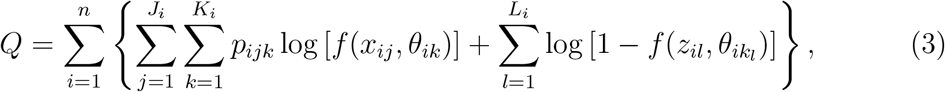

which is exactly a binary cross-entropy loss function, commonly used in deep learning, except with weight *p*_*ijk*_ for binders and weight 1 for non-binders. For the *j*-th binder from sample *i*, we duplicate it *K*_*i*_ times and assign the weights *p*_*ijk*_ to them, and *f*(*, *θ*_*ik*_ can then be estimated by training the neural network with weighted observations.

### 2.4 Training the neural network

#### 2.4.1 Data Encoding

The first step of the neural network training is to determine how the peptides are encoded as input to the neural network. Most peptides presented by HLA-I proteins are 8-11 AA long, with the majority being 9 AA and the longest ones being 15 AA long. Some previous methods train one neural network for each peptide length [34]. However, it is desirable to borrow information across peptides of different lengths since they share similar sequence patterns. This can be done by transforming the peptides to the same length. There are two popular approaches. One is to introduce insertions or deletions of contiguous positions to transform all peptides to 9 AA long, as implemented by NetMHCpan4.1 and some earlier versions of the NetMHC methods [11, 12]. The other is to transform all peptides to a 15 AA representation, while preserving the four AAs at both ends and the AA(s) in the middle. Other positions are filled with the wild card “X”. This approach is used in MHCflurry [9] and the conservation of AAs at both ends are supported by findings of earlier works [34]. More details of the peptide length transformation are presented in the Supplementary Materials Section B.1. We implemented both approaches of peptide length transformation and concluded that 15 AA encoding lead to better performance.

We adopted a variation of one-hot encoding to encode amino acids. Specifically, we encoded each amino acid by a vector of length 21. Each position of this vector corresponded to one of the twenty amino acids and the wild card “X”. Following earlier work of NetMHC [11], each element of this vector was encoded as 0.05 except the position corresponding to the observed amino acid, which was set to be 0.9. A peptide sequence of length *m* would be encoded to a numerical matrix of size *m* × 21 and further flattened to a vector of length 21*m* as part of the input to the neural network. Using the 15-length representation for a peptide, the peptide input encoding would be a vector of length 315 = 15 × 21. The HLA sequence was represented by 34 positions with close contact to peptides [12]. This 34 AA pseudo-sequence was similarly encoded as a vector of length 34 × 21 = 714. Using the 15 AA representation of peptide sequence and the 34 AA pseudo-sequence for HLA, the final size of the input to the neural network was a vector of length 1029 =(34 + 15) × 21 (Figure 1 (B)).

#### 2.4.2 Neural network design and training

The training data were separated into 5 splits [32]. We used one split as training and another split as validation to explore different architectures of the neural network or the parameters of model training. None of the testing datasets were used to select the final version of the neural networks in PEPPRMINT.

We considered neural networks with one or two hidden dense layers with rectified linear unit (relu) activation function. For the one layer neural network, we considered a dense layer with 100, 200, 400, or 800 nodes. If there was a second dense layer, the number of nodes was half of the number of nodes in the first layer. Each dense layer was followed by a drop out layer. Finally, the last hidden layer was connected to an output layer with a single output and the sigmoid activation function (Figure 1(B)). From the results in validation data, we chose three architectures: a single layer with 200, 400, and 800 nodes. We also considered alternative architectures, such as applying a convolutional neural network on the peptide encoding, or passing the encoded peptide and HLA to separate dense layers before concatenating them. None of these alternative architectures delivered better performance.

The neural network was trained with stochastic gradient descent (Adam optimizer, batch size 32, learning rate 0.001) and with a drop-out rate of 0.5. It was first trained using SA data. Then, using those results as initial values, we trained the mixture model up to 10 EM iterations with the MA data. In each EM iteration, the neural network was trained for 10 epochs. Each peptide was assigned a weight derived from the previous EM iteration. Due to the stochastic property of the neural network, it was possible that the log likelihood in one EM iteration may be lower than the one in the previous EM iteration. Thus, model training was stopped earlier if the log likelihood converged or decreased more than twice and the previous model was saved as the final model. We explored other model training options, e.g., adding *l*_1_/*l*_2_ regularization, adjusting the hyper-parameters (learning rate, batch size, drop out rate, number of epochs and iterations, etc.), assigning class weight for binders vs. non-binders, and the results of other options in the validation data were either similar or worse to the current option.

Another consideration was that the initial neural network estimated by the SA data might prefer to assign peptides to the HLAs included in the SA data, which might lead to over-estimation of the mixture proportions of those HLAs and re-enforce such bias. To avoid this situation, we slowed down the estimate of mixture proportions by replacing the MLE of the mixture proportions in the current iteration with the average of the mixture proportions of the previous iteration and the current iteration.

### 2.5 Evaluation of PEPPRMINT and other methods

We evaluated the performance of PEPPRMINT as well as two existing methods, NetMHCpan-4.1 [32] and MHCflurry-2.0 [16], using the aforementioned MA test set and SA test set. When making prediction for a peptide in the MA test set by a specific neural network, we took the maximum score across all possible HLAs as the output of this neural network. Altogether 15 neural networks were included in PEPPRMINT for three model configurations (single layer with 200, 400, and 800 nodes) and five splits of the training data. We aggregated the results of these 15 neural networks by taking the average of the maximum scores. For both NetMHCpan-4.1 and MHCflurry 2.0, we used the maximum score for each peptide across all possible HLAs for evaluations. Method performance was evaluated using area under the receiver operating characteristic curve (AUC), as well as positive predictive value (PPV). PPV for a sample was calculated by ordering all peptides by their prediction and finding the proportion of binders in the top *N* peptides, where *N* was the total number of true binders for that sample.

## 3 Method Evaluation using Testing Data

### 3.1 Aggregation of the Results of Multiple Neural Networks Improves Performance

The performance across the 15 neural networks trained by PEPPRMINT were similar in both the MA test set and the SA test set as shown in Table 1. Aggregated results led to better performance compared to any of the individual neural networks in the MA test set and better performance than most of the individual neural networks in the SA test set, supporting the aggregation of the 15 neural networks.

**Table 1:** AUC performance across the 15 models and aggregated score. All hyperparameters of the models were the same, except for the number of hidden nodes in the neural network (200, 400, or 800). The training data was divided into 5 splits and each neural network is trained by one split of the data.

| Nodes, Train Data | MA Test Set | SA Test Set |
| --- | --- | --- |
| 200, split 0 | 0.905 | 0.912 |
| 200, split 1 | 0.907 | 0.914 |
| 200, split 2 | 0.903 | 0.901 |
| 200, split 3 | 0.905 | 0.906 |
| 200, split 4 | 0.902 | 0.903 |
| 400, split 0 | 0.905 | 0.913 |
| 400, split 1 | 0.903 | 0.912 |
| 400, split 2 | 0.905 | 0.899 |
| 400, split 3 | 0.901 | 0.900 |
| 400, split 4 | 0.901 | 0.896 |
| 800, split 0 | 0.904 | 0.915 |
| 800, split 1 | 0.902 | 0.906 |
| 800, split 2 | 0.908 | 0.897 |
| 800, split 3 | 0.904 | 0.904 |
| 800, split 4 | 0.905 | 0.907 |
| Aggregated | 0.909 | 0.913 |

### 3.2 Comparison to other methods

Other than PEPPRMINT, NetMHCpan-4.1 is the only other neural network method that handles MA training data. In the MA test set, PEPPRMINT had a higher overall AUC and PPV than NetMHCpan-4.1 (0.909 vs. 0.877 and 0.368 vs. 0.339, respectively). When evaluating the performance for each sample, PEPPRMINT out-performed NetMHCpan-4.1 in 13 of 20 samples (Figure 3(A)). Compared with MHCflurry-2.0, which was trained using SA data, PEPPRMINT performed better or equivalent in 19 of the 20 samples (Figure 3(B)). KESKIN 13240-005 was the only sample where PEPPRMINT’s performance was worse than MHCflurry-2.0. This sample had 472 binders, which was relatively small compared to other samples, of which the median number of binders was 1032. We also examined the sample-by-sample PPVs; PEPPRMINT had similar performance to NetMHpan-4.1 and performed better than or similar to MHCflurry-2.0 in 19 out of 20 samples. More details of the results are summarized in the Supplementary Materials Section B.3.

**Figure 3:**
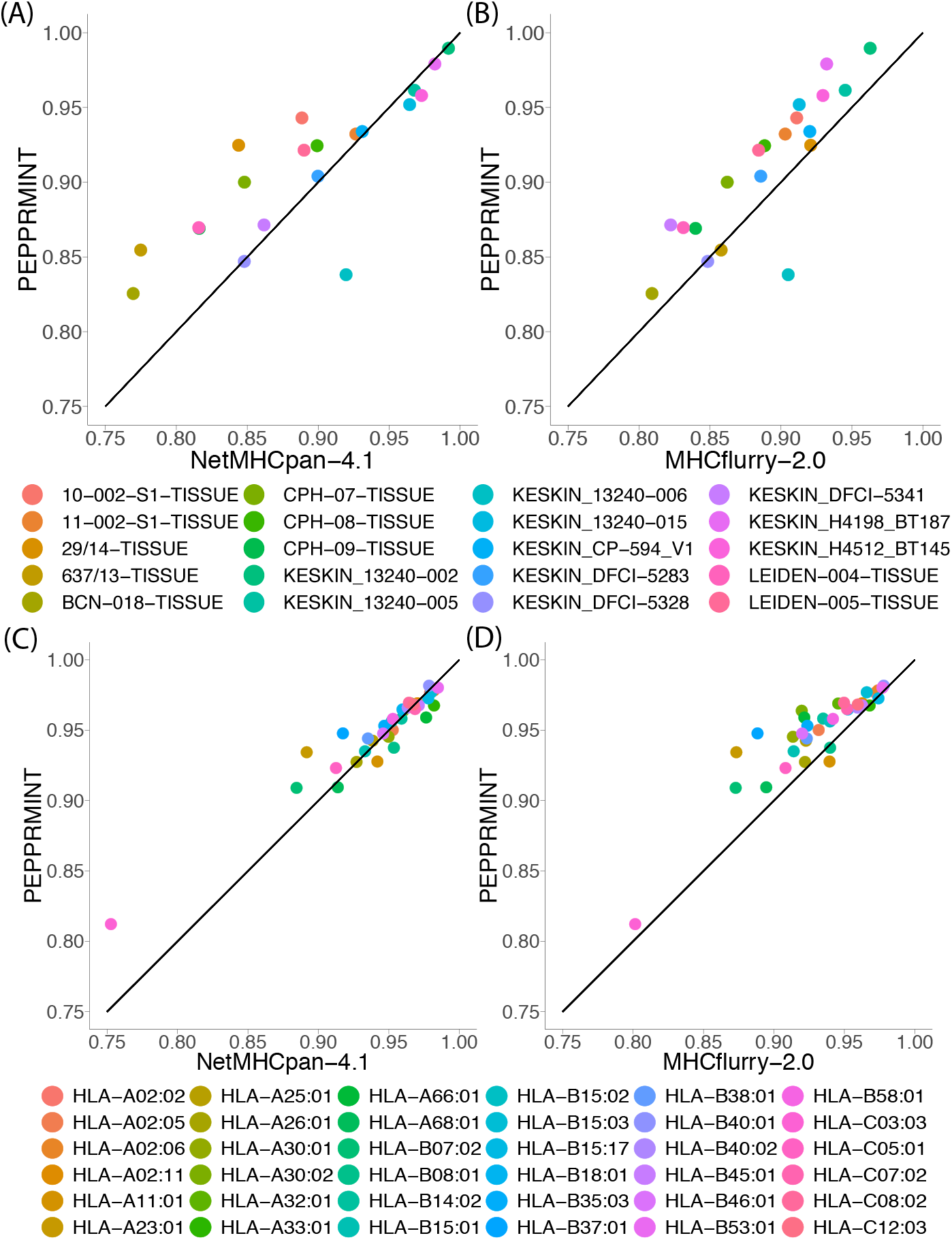
AUC performance comparison. Each point is a sample (for the MA test set) or an HLA (for the SA test set). Points above the diagonal indicate PEPPRMINT has better prediction performance. (A) NetMHCpan-4.1 vs. PEPPRMINT in the MA test set. (B) MHCflurry-2.0 vs. PEPPRMINT in the MA test set. (C) NetMHCpan-4.1 vs. PEPPRMINT in the SA test set. (D) MHCflurry 2.0 vs. PEPPRMINT in SA test set.

The SA test set was used as a secondary benchmark to further evaluate our method. PEPPRMINT performed similar to NetMHCpan-4.1 in most of the HLAs and had a more apparent advantage than NetMHCpan-4.1 in 4 out of the 36 HLAs (Figure 3(C)). PEPPRMINT out performed MHCflurry-2.0 in 34 out of the 36 HLAs (Figure 3(D)).

## 4 Prioritizing Candidates for Cancer Vaccine for Melanoma Patients

We analyzed the whole exome sequence data (around 150x coverage) from 90 tumor samples with paired normal samples from 60 melanoma patients who were treated by Nivolumab, a check-point inhibitor that targets PD-1 [24]. Among the 90 tumor samples, 59 were from pre-therapy biopsies and 31 were from on-therapy biopsies. We first applied our pipeline to call and extract peptide sequences around the somatic mutations, either the mutated peptides or wild-type peptides. More details are supplied in the Supplementary Materials Section E. The number of non-synonymous single nucleotide variants (SNVs) varied from 1 to more than 6,000 (Supplementary Figure 1), with a median number of somatic mutations being 179 and 149 for the pre-therapy or the on-therapy samples, respectively.

Since our evaluations in the previous section showed that PEPPRMINT outperformed MHCflurry-2.0 in most cases, we only compared PEPPRMINT and NetMHCpan-4.1 on this melanoma dataset. For each SNV in a tumor sample, we considered all overlapping 9 AA mutated peptides and all possible HLAs to find the maximum association score of these peptide-HLA pairs, referred to as the neoantigen score for this somatic mutation. Analogously, we calculated a neoantigen score using 9 AA wild type peptides. See Supplementary Materials Section C for more details.

We found the neoantigen scores were strongly correlated between the mutated and reference peptides using either the estimates by NetMHCpan-4.1 or PEPPRMINT as shown in Figure 4 (A)-(B). However, the neoantigen scores calculated by NetMHCpan-4.1 tended to be smaller and had larger divergence between the reference peptides and mutated peptides (Figure 4(A)).

**Figure 4:**
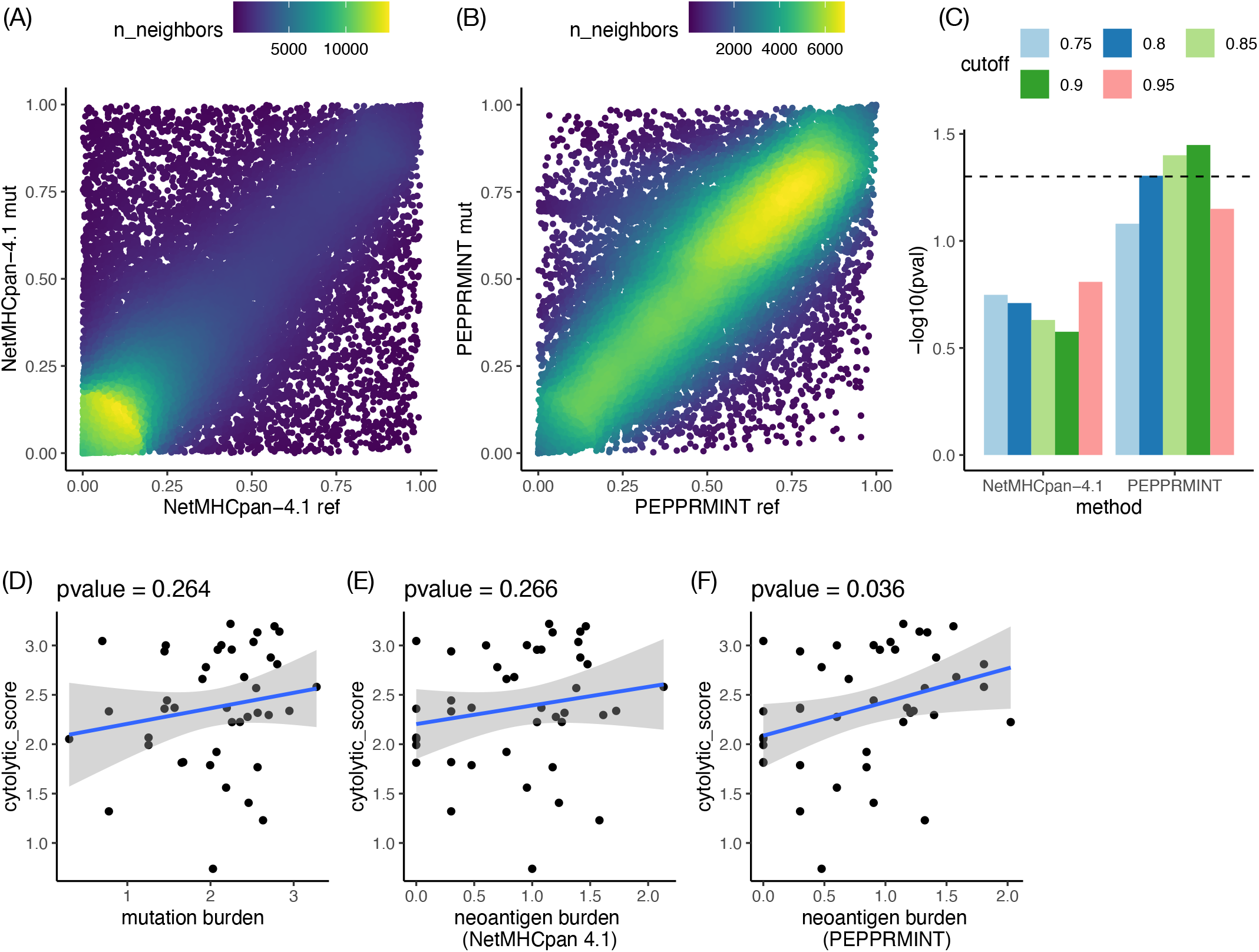
(A-B) Neoantigen scores for 31,298 somatic SNVs calculated by NetMHCpan-4.1 and PEPPRMINT. (C) Summary of −log_10_(p-value) of the associations between neoantigen burdens and cytolytic score. (D-F) Scatter plot of cytolytic score vs mutation burden or neoantigen burden with score cutoff = 0.9 (NetMHCpan 4.1 or PEPPRMINT). Blue line is a linear fit line and the p-value on top of each panel is the association p-value.

While it was difficult to assess whether each neoantigen could induce immune response without additional experiments, it can be indirectly measured by the collective effect of the somatic mutations with high neoantigen scores. More specifically, we defined neoantigen burden as the log10-transformed total number of somatic mutations with neoantigen scores larger than one of a few cutoffs: 0.75, 0.8, 0.85, 0.9, and 0.95. Then, we checked whether neoantigen burden (pre-therapy) was associated with cytolytic score, a measurement of immune system activity (pre-therapy) using the gene expression of immune related genes [24, 35]. We conducted our analysis for 41 pre-therapy samples samples with gene expression available. The neoantigen burdens derived from PEPPRMINT consistently showed stronger associations than the neoantigen burdens from NetMHCpan-4.1 (Figure 4 (C,E,F)). Mutation burden, defined as the log10-transformed total number of non-synonymous SNVs per sample and commonly used as an predictor for immunotherapy response, also had an insignificant association with cytolytic score (Figure 4 (D)).

Among the patients with both pre-therapy and on-therapy samples, patients who responded to the anti-PD1 treatment (partial response or complete response, PRCR) tended to have a larger reduction of neoantigen burdens than non-responding patients (progressive disease or stable disease, PDSD) (Supplementary Figure 3). This suggested that immunotherapy could not invigorate immune system in those nonresponders, even though some of them had a larger number of neoantigens. Therefore, if a cancer vaccine could help the immune system recognize some of these neoantigens, then combining cancer vaccines and checkpoint inhibitors could improve the treatment response in this patient population.

A wild-type peptide is unlikely to induce a strong immune response even if it is presented at the cell surface; however, a strong similarity between the wild-type and the mutated peptide might cause a reduction in the immune response to the mutated peptide. Thus, we were interested in the neoantigens where the mutated peptides around the somatic mutations had high neoantigen scores and the wild-type peptides had low neoantigen scores, so that the potential immune response could be more specific to tumor cells. We defined a high neoantigen score for mutated peptides as > 0.9 and the threshold for the wild type peptides with a low neoantigen score was defined as the median across all somatic mutations: 0.5 for PEPPRMINT and 0.22 for NetMHCpan-4.1. Using this approach, we selected 141 and 148 neoantigens in 39 and 30 patients by NetMHCpan-4.1 and PEPPRMINT, respectively (Supplementary Files 2 and 3). Interestingly, the patients who responded to anti-PD1 treatment were more likely to have at least one such neoantigen. For NetMHCpan-4.1, 12 of 13 responders and 27 out of 47 non-responders had at least one such neoantigen. For PEPPRMINT, 10 out of 13 responders and 20 out of 47 non-responders had at least one neoantigen.

## 5 Discussion

Cancer immunotherapy, particularly immune checkpoint inhibitors, have achieved impressive successes in cancer treatment. However, durable clinical benefits are only observed in a subset of patients [36]. Cancer vaccine development is a promising approach to expand the patient population that could benefit from immunotherapy [37]. Since only a very small proportion of peptides are presented on the cell surface, computational prioritization of peptides presented by HLA-I proteins on the cell surface is a very important step for cancer vaccine design. To successfully prioritize peptides, one promising way is to develop efficient models for MA data, since such data are more accurate and expected to accumulate quickly in the near future.

Currently, the only other method that is able to model MA data by neural networks is NetMHCpan-4.1 [32]. In this paper, we proposed an alternative method called PEPPRMINT, which combines a rigorous statistical model (a mixture model) and the power of a neural network to analyze MA data. PEPPRMINT performed better or similarly to NetMHCpan-4.1. However, since the mixture mode framework is more flexible to model MA data, we expect the performance of PEPPRMINT to improve faster than NetMHCpan-4.1 as these models are updated using more training data. Furthermore, application of PEPPRMINT and NetMHCpan-4.1 in a case study to prioritize neoantigens for melanoma patients demonstrated that the neoantigen burden calculated by PEPPRMINT had a significant association with cytolytic activity, while NetMHCpan-4.1 did not capture this association. Our method is computationally efficient to make predictions. For example, in our melanoma application, it took about 2 minutes to make prediction for around 280,000 peptides using one neural network.

There are several directions for future studies to improve our pipeline. First, we may include the 3-5 amino acids flanking a peptide into our model, since earlier studies have shown that these locations can provide information regarding to peptide processing [34, 38]. It is expected that information in flanking regions are the same across HLAs since peptide processing is independent of HLAs. Therefore, the flanking sequences could be modeled outside the mixture model to reduce model complexity and improve computational efficiency. Another direction is to incorporate gene expression information to prioritize neoantigens. The sequence-based prediction and gene expression could be combined through a simple regression model or another neural network [34, 38]. Additionally, in this study, we focused on SNVs. In the future, expansion to other types of somatic mutations, such as indels or splicing variants, would be desirable.

Finally, while our work focuses on HLA-I neoantigens, another important future direction is to prioritize neoantigens for HLA-II proteins. HLA-I and HLA-II proteins present antigens to CD8+ and CD4+ T cells, respectively. Emerging evidence has demonstrated the importance of HLA-II neoantigens and CD4+ T cells in eliciting response to immune checkpoint inhibitors [39–41]. For example, Alspach et al. [40] showed that neoantigen-specific CD8+ and CD4+ T cells are both required for immunotherapy induced anti-tumor response. Cancer vaccine studies have also shown that HLA-II neoantigens elicit CD4+T cell response and confer strong anti-tumour activity [2, 42, 43].

## Supporting information

Supplementary Materials for "PEPPRMINT: a statistical model to predict peptide presentation by HLA-I proteins"

## 6 Code availability

PEPPRMINT source code, data, and trained models are available at https://github.com/Sun-lab/PEPPRMINT. Pipelines for neoantigen prioritization and code for Riaz et al data analysis are available at https://github.com/Sun-lab/IT-predictor.

## References

[1] Finck, A., Gill, S. I., and June, C. H. (2020) Cancer immunotherapy comes of age and looks for maturity. Nature Communications, 11(1), 1–4.

[2] Ott, P. A., Hu, Z., Keskin, D. B., Shukla, S. A., Sun, J., Bozym, D. J., Zhang, W., Luoma, A., Giobbie-Hurder, A., Peter, L., et al. (2017) An immunogenic personal neoantigen vaccine for patients with melanoma. Nature, 547(7662), 217–221.

[3] Keskin, D. B., Anandappa, A. J., Sun, J., Tirosh, I., Mathewson, N. D., Li, S., Oliveira, G., Giobbie-Hurder, A., Felt, K., Gjini, E., et al. (2019) Neoantigen vaccine generates intratumoral T cell responses in phase Ib glioblastoma trial. Nature, 565(7738), 234–239.

[4] Hu, Z., Leet, D. E., Allesøe, R. L., Oliveira, G., Li, S., Luoma, A. M., Liu, J., Forman, J., Huang, T., Iorgulescu, J. B., et al. (2021) Personal neoantigen vaccines induce persistent memory T cell responses and epitope spreading in patients with melanoma. Nature medicine, 27(3), 515–525.

[5] Peng, M., Mo, Y., Wang, Y., Wu, P., Zhang, Y., Xiong, F., Guo, C., Wu, X., Li, Y., Li, X., et al. (2019) Neoantigen vaccine: an emerging tumor immunotherapy. Molecular cancer, 18(1), 1–14.

[6] Lundegaard, C., Lamberth, K., Harndahl, M., Buus, S., Lund, O., and Nielsen, M. (2008) NetMHC-3.0: accurate web accessible predictions of human, mouse and monkey MHC class I affinities for peptides of length 8–11. Nucleic acids research, 36(suppl 2), W509–W512.

[7] Kim, Y., Sidney, J., Pinilla, C., Sette, A., and Peters, B. (2009) Derivation of an amino acid similarity matrix for peptide: MHC binding and its application as a Bayesian prior. BMC bioinformatics, 10(1), 1–11.

[8] Nielsen, M. and Andreatta, M. (2016) NetMHCpan-3.0; improved prediction of binding to MHC class I molecules integrating information from multiple receptor and peptide length datasets. Genome medicine, 8(1), 1–9.

[9] O’Donnell, T. J., Rubinsteyn, A., Bonsack, M., Riemer, A. B., Laserson, U., and Hammerbacher, J. (2018) MHCflurry: open-source class I MHC binding affinity prediction. Cell systems, 7(1), 129–132.

[10] Gfeller, D. and Bassani-Sternberg, M. (2018) Predicting antigen presentation–what could we learn from a million peptides?. Frontiers in immunology, 9, 1716.

[11] Andreatta, M. and Nielsen, M. (2016) Gapped sequence alignment using artificial neural networks: application to the MHC class I system. Bioinformatics, 32(4), 511–517.

[12] Jurtz, V., Paul, S., Andreatta, M., Marcatili, P., Peters, B., and Nielsen, M. (2017) NetMHCpan-4.0: improved peptide–MHC class I interaction predictions integrating eluted ligand and peptide binding affinity data. The Journal of Immunology, 199(9), 3360–3368.

[13] Bhattacharya, R., Sivakumar, A., Tokheim, C., Guthrie, V. B., Anagnostou, V., Velculescu, V. E., and Karchi, R. (2017) Evaluation of machine learning methods to predict peptide binding to MHC Class I proteins. bioRxiv,.

[14] Liu, Z., Cui, Y., Xiong, Z., Nasiri, A., Zhang, A., and Hu, J. (2018) DeepSeqPan, a novel deep convolutional neural network model for pan-specific class I HLA-peptide binding affinity prediction. bioRxiv, p. 299412.

[15] Robinson, J., Halliwell, J., Hayhurst, J., Flicek, P., Parham, P., and Marsh, S. (2015) The IPD and IMGT/HLA database: allele variant databases. Nucleic acids research, 43(Database issue), D423–31.

[16] O’Donnell, T. J., Rubinsteyn, A., and Laserson, U. (2020) MHCflurry 2.0: Improved pan-allele prediction of MHC class I-presented peptides by incorporating antigen processing. Cell systems, 11(1), 42–48.

[17] Trowsdale, J. and Knight, J. C. (2013) Major histocompatibility complex genomics and human disease. Annual review of genomics and human genetics, 14, 301–323.

[18] Bassani-Sternberg, M., Chong, C., Guillaume, P., Solleder, M., Pak, H., Gannon, P. O., Kandalaft, L. E., Coukos, G., and Gfeller, D. (2017) Deciphering HLA-I motifs across HLA peptidomes improves neo-antigen predictions and identifies allostery regulating HLA specificity. PLoS computational biology, 13(8), e1005725.

[19] Alvarez, B., Reynisson, B., Barra, C., Buus, S., Ternette, N., Connelley, T., Andreatta, M., and Nielsen, M. (2019) NNAlign MA; MHC peptidome deconvolution for accurate MHC binding motif characterization and improved T cell epitope predictions. Molecular & Cellular Proteomics, p. in press.

[20] Bassani-Sternberg, M. and Gfeller, D. (2016) Unsupervised HLA peptidome deconvolution improves ligand prediction accuracy and predicts cooperative effects in peptide–HLA interactions. The Journal of Immunology, 197(6), 2492–2499.

[21] Racle, J., Michaux, J., Rockinger, G., Arnaud, M., Bobisse, S., Chong, C., Guillaume, P., Coukos, G., Harari, A., Jandus, C., Bassani-Sternberg, M., and Gfeller, D. (2019) Robust prediction of HLA class II epitopes by deep motif deconvolution of immunopeptidomes. Nature Biotechnology, p. in press.

[22] Andreatta, M., Lund, O., and Nielsen, M. (2012) Simultaneous alignment and clustering of peptide data using a Gibbs sampling approach. Bioinformatics, 29(1), 8–14.

[23] Sade-Feldman, M., Jiao, Y. J., Chen, J. H., Rooney, M. S., Barzily-Rokni, M., Eliane, J.-P., Bjorgaard, S. L., Hammond, M. R., Vitzthum, H., Blackmon, S. M., et al. (2017) Resistance to checkpoint blockade therapy through inactivation of antigen presentation. Nature communications, 8(1), 1–11.

[24] Riaz, N., Havel, J. J., Makarov, V., Desrichard, A., Urba, W. J., Sims, J. S., Hodi, F. S., Martín-Algarra, S., Mandal, R., Sharfman, W. H., et al. (2017) Tumor and microenvironment evolution during immunotherapy with nivolumab. Cell, 171(4), 934–949.

[25] Li, H. and Durbin, R. (2009) Fast and accurate short read alignment with Burrows–Wheeler transform. bioinformatics, 25(14), 1754–1760.

[26] Van der Auwera, G. A. and O’Connor, B. D. (2020) Genomics in the Cloud: Using Docker, GATK, and WDL in Terra, O’Reilly Media, .

[27] Cibulskis, K., Lawrence, M. S., Carter, S. L., Sivachenko, A., Jaffe, D., Sougnez, C., Gabriel, S., Meyerson, M., Lander, E. S., and Getz, G. (2013) Sensitive detection of somatic point mutations in impure and heterogeneous cancer samples. Nature biotechnology, 31(3), 213–219.

[28] Saunders, C. T., Wong, W. S., Swamy, S., Becq, J., Murray, L. J., and Cheetham, R. K. (2012) Strelka: accurate somatic small-variant calling from sequenced tumor–normal sample pairs. Bioinformatics, 28(14), 1811–1817.

[29] Wang, K., Li, M., and Hakonarson, H. (2010) ANNOVAR: Functional annotation of genetic variants from next-generation sequencing data. Nucleic Acids Research, 38, e164.

[30] Szolek, A., Schubert, B., Mohr, C., Sturm, M., Feldhahn, M., and Kohlbacher, O. (2014) OptiType: precision HLA typing from next-generation sequencing data. Bioinformatics, 30(23), 3310–3316.

[31] Nielsen, M., Lundegaard, C., Blicher, T., Lamberth, K., Harndahl, M., Justesen, S., Røder, G., Peters, B., Sette, A., Lund, O., et al. (2007) NetMHCpan, a method for quantitative predictions of peptide binding to any HLA-A and-B locus protein of known sequence. PloS one, 2(8), e796.

[32] Reynisson, B., Alvarez, B., Paul, S., Peters, B., and Nielsen, M. (2020) NetMHCpan-4.1 and NetMHCIIpan-4.0: improved predictions of MHC antigen presentation by concurrent motif deconvolution and integration of MS MHC eluted ligand data. Nucleic Acids Research,.

[33] Sarkizova, S., Klaeger, S., Le, P. M., Li, L. W., Oliveira, G., Keshishian, H., Hartigan, C. R., Zhang, W., Braun, D. A., Ligon, K. L., et al. (2020) A large peptidome dataset improves HLA class I epitope prediction across most of the human population. Nature biotechnology, 38(2), 199–209.

[34] Bulik-Sullivan, B., Busby, J., Palmer, C. D., Davis, M. J., Murphy, T., Clark, A., Busby, M., Duke, F., Yang, A., Young, L., et al. (2019) Deep learning using tumor HLA peptide mass spectrometry datasets improves neoantigen identification. Nature biotechnology, 37(1), 55.

[35] Rooney, M. S., Shukla, S. A., Wu, C. J., Getz, G., and Hacohen, N. (2015) Molecular and genetic properties of tumors associated with local immune cytolytic activity. Cell, 160(1-2), 48–61.

[36] Havel, J. J., Chowell, D., and Chan, T. A. (2019) The evolving landscape of biomarkers for checkpoint inhibitor immunotherapy. Nature Reviews Cancer, 19(3), 133–150.

[37] Blass, E. and Ott, P. A. (2021) Advances in the development of personalized neoantigen-based therapeutic cancer vaccines. Nature Reviews Clinical Oncology, 18(4), 215–229.

[38] Chen, B., Khodadoust, M. S., Olsson, N., Wagar, L. E., Fast, E., Liu, C. L., Muftuoglu, Y., Sworder, B. J., Diehn, M., Levy, R., et al. (2019) Predicting HLA class II antigen presentation through integrated deep learning. Nature biotechnology, 37, 1332–1343.

[39] Borst, J., Ahrends, T., Babala, N., Melief, C. J., and Kastenmuller, W. (2018) CD4+ T cell help in cancer immunology and immunotherapy. Nature Reviews Immunology, 18(10), 635–638.

[40] Alspach, E., Lussier, D. M., Miceli, A. P., Kizhvatov, I., DuPage, M., Luoma, A. M., Meng, W., Lichti, C. F., Esaulova, E., Vomund, A. N., et al. (2019) MHC-II neoantigens shape tumour immunity and response to immunotherapy. Nature, 574(7780), 696–701.

[41] Tay, R. E., Richardson, E. K., and Toh, H. C. (2020) Revisiting the role of CD4+ T cells in cancer immunotherapy—new insights into old paradigms. Cancer Gene Therapy, pp. 1–13.

[42] Kreiter, S., Vormehr, M., Van de Roemer, N., Diken, M., Löwer, M., Diekmann, J., Boegel, S., Schrörs, B., Vascotto, F., Castle, J. C., et al. (2015) Mutant MHC class II epitopes drive therapeutic immune responses to cancer. Nature, 520(7549), 692–696.

[43] Sahin, U., Derhovanessian, E., Miller, M., Kloke, B.-P., Simon, P., Löwer, M., Bukur, V., Tadmor, A. D., Luxemburger, U., Schrörs, B., et al. (2017) Personalized RNA mutanome vaccines mobilize poly-specific therapeutic immunity against cancer. Nature, 547(7662), 222–226.

