## Supplementary Materials for "PEPPRMINT: a statistical model to predict peptide presentation by HLA-I proteins" for "Prioritizing candidate peptides for cancer vaccines by PEPPRMINT: a statistical model to predict peptide presentation by HLA-I proteins": Supplementary_PEPPRMINT.pdf

### A Existing Methods to Predict Peptide Presentation by HLAs

#### A.1 Introduction

Earlier methods aimed at predicting peptide presentation by HLAs used unsupervised clustering of peptides through mixture models (Bassani-Sternberg and Gfeller, 2016; Racle et al., 2019) and Gibbs sampler (Andreatta et al., 2013, 2017). These methods quantified the likelihood of peptide presentation using a position weight matrix (PWM), which described the marginal distribution of AA at each position of the peptide and ignored their interactions. Furthermore, these unsupervised methods may not have fully recovered all clusters because different HLA alleles may have similar binding motifs or due to the very low expression of some HLA alleles (Bassani-Sternberg and Gfeller, 2016).

Neural network (NN) methods have been shown to make more accurate peptide presentation prediction than PWM-based methods, likely due to their capability to take account of the interactions across AAs (Zhao and Sher, 2018), such as Bulik-Sullivan et al. (2019), MHCflurry-2.0 (O'Donnell et al., 2020), and NetMHC-Pan/NNAlign methods (Nielsen and Andreatta (2016); Reynisson et al. (2020)). Bulik-Sullivan et al. (2019) developed a deep learning method that predicts peptide presentation for each HLA-I allele separately using MS data. However, this method did not borrow information across HLA alleles and was unable to make predictions on HLA alleles with insufficient training data. NetMHC and NNAlign methods are further described in the next section.

#### A.2 NNAlign, NNAlign_MA, and NetMHCpan-4.1

NNAlign is an HLA binding affinity prediction method trained on single allele (SA) binding affinity and eluted ligand data that borrows information across alleles to increase the predictive power (Nielsen and Andreatta, 2017). NNAlign uses artificial neural networks to identify linear motifs in biological sequences and generate a model of the sequence motif detected in the data. It employs the pan-specific approach, i.e. each HLA allele is represented by its sequence, and thus it can be used to make prediction for any HLA allele with sequence information. More specifically, following some earlier works of NetMHC methods, it uses a pseudo-sequence of an HLA-I allele derived from the Anthony Nolan database instead of the whole HLA sequence as input to the model (Nielsen et al., 2007). This method reduces the full HLA sequence to only 34 AA, as shown in Figure 4 of Nielsen et al. (2007).

Alvarez et al. (2019) extends NNAlign to multi-allele (MA) data and proposes NNAlign_MA, a pan-specific method that takes both the peptide and the HLA sequence as input to predict peptide presentation for any HLA-I alleles. NNAlign_MA fully automates clustering and labeling of mass spectrum eluted ligand data. NNAlign_MA uses an integrated framework to iteratively update the clustering, MHC (Major Histocompatibility Complex) annotation, and peptide binding predictions. Three types of training data are used: SA binding affinity data, SA eluted ligand data, and MA eluted ligand data. All training data is supplemented with negative peptides by extracting random peptides from the UniProt database. For MA data, a random HLA from the corresponding sample is assigned to each negative peptide.

NNAlign_MA is first trained by the SA data. Then the model is iteratively trained using the MA data. In each iteration, each peptide in the MA dataset is predicted for all possible HLAs in the corresponding sample and the peptide is "annotated" by the HLA allele with the highest predictive value. This annotation procedure transforms the MA data to SA data. The SA data and the annotated MA data are merged and the model is re-trained on the combined data. Models are trained for 200 iterations. NNAlign_MA trains multiple neural networks. All networks have a single hidden layer and an output layer with two values, one for binding affinity and one for the eluted ligand. The training data is split into five folds. The final ensemble of models includes 250 networks (2, 10, 20, 40 and 60 hidden neurons and 10 random weight initiation seeds for each fold). NetMHCpan-4.1 is the collection of neural networks of NNAlign_MA trained by a recent dataset (Reynisson et al., 2020).

As input to the NetMHCpan-4.1 or NNAlign_MA, peptides are processed to be 9 AA in length. If the peptide is longer than 9 AA, all possible consecutive deletions are applied to the sequence to determine a core of 9 AA and the sequence with the strongest affinity is selected. If the peptide is 8 AA, then a wildcard AA 'X' is added to each possible position of the peptide. Peptides are encoded as a 20-digit binary number for each AA, e.g., 0.90 if that position has that AA otherwise 0.05 or Blosum50 (Henikoff and Henikoff (1992)). Affinities are capped at 50,000 and then transformed as $1 - \log(a)/\log(50000)$, where $a$ = affinity, to output a value between 0 and 1. In addition to the peptide, NetMHCpan-4.1 also takes the 34 AA pseudo-sequence of the HLA-I allele as part of the input.

##### A.3 MHCflurry and MHCflurry-2.0

MHCflurry is a feed forward HLA-I allele-specific NN method trained on binding affinity and SA data (O'Donnell et al., 2018). The NN models contain zero to two locally connected layers with no weight sharing, a fully connected layer, and an output layer with sigmoid activation function. Of the 320 models trained per allele, 8-16 models are selected using forward step-wise selection. MHCflurry uses a novel fixed-length encoding that preserves the positionality of the AA that have important stabilizing contacts with the HLA. It is commonly believed that the "anchor positions" of peptides, or positions that are more important to binding to MHC, are at the beginning or the end of the peptide. MHCflurry encodes all 8-15 length peptides as a 15 length sequence, where missing AA are filled in with "X". See Section B.1 for more details. Though, MHCflurry can only predict for HLA alleles that are in the training data, the 15-length representation utilizes more information than the NetMHC methods since it does not cut longer peptides to 9 AA.

MHCflurry-2.0 is a pan-allelic method that builds upon MHCflurry by combining binding affinity and antigen processing (AP) signals O'Donnell et al. (2020). First, a binding affinity predictor is trained with binding affinity and mass spectrum SA data. A training set of AP signals is identified by selecting those binders and non-binders with high affinity prediction. The AP predictor models the residual allele-independent sequence properties that was not learned in the binding affinity predictor. MHCflurry-2.0 uses a 45-length representation, by concatenating the left aligned, centered, and right aligned 15-length representations with "X" filled in the missing spaces. For example, the peptide "AMDGILGFV" is represented by concatenating "AMDGILGFVXXXXXX" (left aligned), "XXXAMDGILGFVXXX" (centered), and "XXXXXXAMDGILGFV" (right aligned), resulting in "AMDGILGFVXXXXXXXXXAMDGILGFVXXXXXXXXXAMDGILGFV". The idea is to ensure that the C and N terminals of the peptide are available to the network at a fixed position in the encoding regardless of peptide length. From 140 trained models, 10 models are selected in the final ensemble using a forward step-wise procedure. More details about training and model selection are available in the STAR METHODS of O'Donnell et al. (2020).

### B Additional Details of PEPPRMINT Method and Results

#### B.1 Model Input Details

Training data for PEPPRMINT and the single-allele mass spectrum (SA) test data are provided by NetMHCpan-4.1 (`http://www.cbs.dtu.dk/suppl/immunology/NAR_NetMHCpan_NetMHCIIpan/`). The training data are called NetMHCpan_train.tar.gz and the SA test data are sorted by HLA. The engine and the downloadable version to run NetMHCpan-4.1 is located at `http://www.cbs.dtu.dk/services/NetMHCpan/`.

The multi-allele mass spectrum (MA) testing data, named Data_S1.csv.gz, are provided by MHCflurry-2.0 at `https://data.mendeley.com/datasets/zx3kjzc3yx/3` (O'Donnell et al., 2020). We use the MULTIALLELIC-RECENT dataset. There are no overlaping samples in the MULTIALLELIC-RECENT and the NetMHCpan-4.1 training data, though there are some overlaping peptides. This is to be expected as there are overlapping peptides across HLA alleles of the SA dataset as well.

For peptide encoding, we adopted the 15-length representation used by MHCflurry (O'Donnell et al., 2018). It did not appear that the 45-length representation used by MHCflurry-2.0 had much improvement in performance compared to the 15-length representation. Additionally, the 45-length representation would triple the memory usage and thus make it computationally much harder to train the model. In the 15-length representation, the first four and last four AAs of the peptide mapped to the first four and last four positions. The remaining AA of the peptides were filled in starting from the center of the 15-length representation. For example, the following peptides AMFKLSTH, SEHLKMFAA, and PHALSKPMBS are encoded as AMFKXXXXXXXLSTH, SEHLXXXKXXXMFAA, and PHALXXXSXXKPMBS, respectively.

##### B.2 Configuration Details and Results

We explored different NN structures, including a deeper NN with two layers, separate processing of peptide and HLA as two inputs by dense layer convolutional layers before concatenating them, and using a mixture model to determine the best way to pad or cut different peptide lengths to 9 AAs. Different configurations were trained with one split of the training data and evaluated on another split of the training data as the validation set. These configurations all had poorer performance than the current NN (with one dense layer) used in PEPPRMINT. Within the current structure, we did a grid search to determine the best values for the hyper-parameters, including for batch size ($b = 16, 32, 64$), number of training epochs ($epoch = 4, 10, 25, 50$), the learning rate ($lr = 0.001, 0.0001$), regularization ($l_1$ and $l_2$ penalty combination), and the number of hidden nodes in the NN (nodes = 100, 200, 400, 800, 1600). Additionally, we examined updating the estimate of of the mixture proportions as a function of the current estimate and the previous iterations estimate ($w * \pi_0 + (1 - w) * \pi_1$), where $w = 1, 0.5, 0.1$ and $\pi_0, \pi_1$ were the previous and current iteration estimate, respectively. The NN performance was not sensitive to the changes of these hyper-parameters. These configurations are available through the GitHub repository PEPPRMINT (https://github.com/Sun-lab/PEPPRMINT).

##### B.3 Sample and HLA Specific Results

Supplementary Figure 1 plots the overall AUC for PEPPRMINT and NetMHCpan-4.1 in the MA test set. Supplementary Table 1 summarizes the performance of PEPPRMINT, NetMHCpan-4.1, and MHCflurry-2.0 on the MA test set by sample. Supplementary Table 2 summarizes the performance for the SA test by HLA.

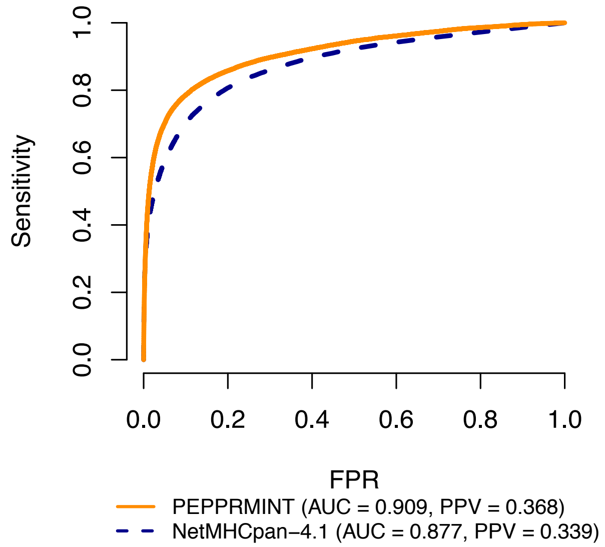

Supplementary Figure 1: Overall AUC performance on MA test set for PEPPRMINT vs. NetMHCPan-4.1

| sample | AUC_PEPPRMINT | AUC_NetMHCpan4.1 | AUC_MHCflurry2 | PPV_PEPPRMINT | PPV_NetMHCpan4.1 | PPV_MHCflurry2 |
|---|---|---|---|---|---|---|
| 10−002−S1−TISSUE | 0.943 | 0.888 | 0.911 | 0.394 | 0.216 | 0.371 |
| 11−002−S1−TISSUE | 0.932 | 0.926 | 0.903 | 0.371 | 0.420 | 0.355 |
| 637/13−TISSUE | 0.855 | 0.775 | 0.858 | 0.212 | 0.068 | 0.325 |
| BCN−018−TISSUE | 0.826 | 0.770 | 0.809 | 0.298 | 0.261 | 0.276 |
| CPH−07−TISSUE | 0.900 | 0.848 | 0.862 | 0.422 | 0.392 | 0.295 |
| CPH−08−TISSUE | 0.924 | 0.899 | 0.888 | 0.430 | 0.432 | 0.315 |
| CPH−09−TISSUE | 0.869 | 0.816 | 0.840 | 0.318 | 0.266 | 0.267 |
| KESKIN_13240−002 | 0.990 | 0.992 | 0.963 | 0.590 | 0.598 | 0.437 |
| KESKIN_13240−005 | 0.962 | 0.968 | 0.945 | 0.461 | 0.617 | 0.424 |
| KESKIN_13240−006 | 0.838 | 0.920 | 0.905 | 0.159 | 0.186 | 0.187 |
| KESKIN_13240−015 | 0.952 | 0.964 | 0.913 | 0.337 | 0.500 | 0.194 |
| KESKIN_CP−594_V1 | 0.934 | 0.931 | 0.920 | 0.396 | 0.521 | 0.328 |
| KESKIN_DFCI−5283 | 0.904 | 0.900 | 0.886 | 0.395 | 0.464 | 0.175 |
| KESKIN_DFCI−5328 | 0.847 | 0.848 | 0.848 | 0.316 | 0.474 | 0.184 |
| KESKIN_DFCI−5341 | 0.871 | 0.862 | 0.822 | 0.210 | 0.328 | 0.175 |
| KESKIN_H4198_BT187 | 0.979 | 0.982 | 0.932 | 0.514 | 0.595 | 0.241 |
| KESKIN_H4512_BT145 | 0.958 | 0.973 | 0.930 | 0.336 | 0.492 | 0.274 |
| LEIDEN−004−TISSUE | 0.870 | 0.816 | 0.831 | 0.312 | 0.248 | 0.272 |
| LEIDEN−005−TISSUE | 0.921 | 0.890 | 0.884 | 0.418 | 0.431 | 0.291 |

Supplementary Table 1: AUC and PPV performance on MA test set for PEPPRMINT, NetMHCPan-4.1, and MHCflurry-2.0 by sample.

| HLA | AUC_PEPPRMINT | PPV_PEPPRMINT | AUC_NetMHCpan4.1 | PPV_NetMHCpan4.1 | AUC_MHCflurry2.0 |
|---|---|---|---|---|---|
| HLA−A02:02 | 0.977 | 0.766 | 0.979 | 0.856 | 0.972 |
| HLA−A02:05 | 0.950 | 0.652 | 0.953 | 0.767 | 0.932 |
| HLA−A02:06 | 0.978 | 0.718 | 0.982 | 0.832 | 0.974 |
| HLA−A02:11 | 0.969 | 0.736 | 0.970 | 0.795 | 0.963 |
| HLA−A11:01 | 0.928 | 0.642 | 0.942 | 0.686 | 0.939 |
| HLA−A23:01 | 0.934 | 0.744 | 0.892 | 0.750 | 0.873 |
| HLA−A25:01 | 0.942 | 0.737 | 0.938 | 0.808 | 0.923 |
| HLA−A26:01 | 0.927 | 0.658 | 0.927 | 0.755 | 0.922 |
| HLA−A30:01 | 0.945 | 0.616 | 0.950 | 0.739 | 0.913 |
| HLA−A30:02 | 0.964 | 0.782 | 0.960 | 0.832 | 0.920 |
| HLA−A32:01 | 0.969 | 0.783 | 0.966 | 0.835 | 0.946 |
| HLA−A33:01 | 0.968 | 0.670 | 0.982 | 0.842 | 0.968 |
| HLA−A66:01 | 0.959 | 0.622 | 0.976 | 0.838 | 0.922 |
| HLA−A68:01 | 0.909 | 0.580 | 0.914 | 0.635 | 0.895 |
| HLA−B07:02 | 0.909 | 0.660 | 0.884 | 0.692 | 0.873 |
| HLA−B08:01 | 0.938 | 0.617 | 0.954 | 0.817 | 0.940 |
| HLA−B14:02 | 0.958 | 0.730 | 0.959 | 0.802 | 0.935 |
| HLA−B15:01 | 0.935 | 0.753 | 0.933 | 0.780 | 0.914 |
| HLA−B15:02 | 0.956 | 0.779 | 0.952 | 0.794 | 0.940 |
| HLA−B15:03 | 0.977 | 0.763 | 0.980 | 0.872 | 0.966 |
| HLA−B15:17 | 0.973 | 0.726 | 0.978 | 0.823 | 0.974 |
| HLA−B18:01 | 0.965 | 0.760 | 0.960 | 0.830 | 0.952 |
| HLA−B35:03 | 0.953 | 0.730 | 0.947 | 0.776 | 0.924 |
| HLA−B37:01 | 0.948 | 0.703 | 0.917 | 0.769 | 0.888 |
| HLA−B38:01 | 0.944 | 0.771 | 0.935 | 0.798 | 0.923 |
| HLA−B40:01 | 0.982 | 0.865 | 0.979 | 0.900 | 0.978 |
| HLA−B40:02 | 0.966 | 0.842 | 0.963 | 0.850 | 0.959 |
| HLA−B45:01 | 0.968 | 0.790 | 0.971 | 0.854 | 0.963 |
| HLA−B46:01 | 0.948 | 0.730 | 0.946 | 0.736 | 0.920 |
| HLA−B53:01 | 0.980 | 0.778 | 0.985 | 0.889 | 0.977 |
| HLA−B58:01 | 0.958 | 0.788 | 0.953 | 0.806 | 0.942 |
| HLA−C03:03 | 0.812 | 0.517 | 0.753 | 0.523 | 0.801 |
| HLA−C05:01 | 0.923 | 0.755 | 0.912 | 0.770 | 0.908 |
| HLA−C07:02 | 0.965 | 0.725 | 0.969 | 0.782 | 0.952 |
| HLA−C08:02 | 0.970 | 0.844 | 0.964 | 0.847 | 0.950 |
| HLA−C12:03 | 0.968 | 0.706 | 0.968 | 0.757 | 0.960 |

Supplementary Table 2: AUC and PPV performance on SA test set for PEPPRMINT, NetMHCPan-4.1, and MHCflurry-2.0 by HLA.

### C Data Processing and Additional Results of Melanoma Data Analysis

#### C.1 Pre-processing and mutation calling

Raw data of FASTQ files were downloaded from the NCBI Sequence Read Archive (SRA, study ID SRP094781) using the `fast_dump` command in the SRA Toolkit. The FASTQ files contained 76 base pair (bp) paired-end whole exome sequencing data. We mapped these sequences to the reference genome (hg38) using Burrows-Wheeler Aligner (BWA) (Li and Durbin, 2009). Before calling mutations, we applied Picard (version 2.20.0) (`http://broadinstitute.github.io/picard/`) to mark duplicated reads and used GATK tools (version 4.1.2.0) to re-calibrate the base quality scores (Van der Auwera and O'Connor, 2020).

Calling somatic mutations is a challenging task. A commonly used strategy to improve the mutation calling accuracy is to combine the results of two or more mutation callers. Based on earlier comparative studies (Xu et al., 2014; Krøigård et al., 2016), we chose two mutation callers: Mutect (v1.1.7) (Cibulskis et al., 2013) and Strelka (v1.0.14) (Saunders et al., 2012) and took the intersection of mutations calls from them. We annotated all the somatic mutations by ANNOVAR (Wang et al., 2010), which provided annotations on the functional impact of a somatic mutation (e.g., whether it alters the protein product), its associated genes, and whether it was observed in the Exome Aggregation Consortium (ExAC) data of germline variants (Lek et al., 2016).

Next, we filtered the somatic mutation calls as follows.

1. Removed all the mutations with coverage less than 20 in either the tumor or paired normal samples.
2. Removed all the mutations with less than 5 reads supporting the alternative allele in the tumor sample.
3. Removed all the mutations with an alternative allele frequency less than 5%.

4. Removed all the mutations with an alternative allele frequency in the Exome Aggregation Consortium (`ExAC_nontcga_ALL` in ANNOVAR output) larger than $10^{-4}$.

Finally, we selected non-synonymous mutations and only kept the single nucleotide variants (SNVs). The number of non-synonymous SNVs per sample varied from 1 to more than 6,723 for the 59 pre-therapy samples, with a median of 179 (Supplementary Figure 1 (A)). It varied from 1 to 957 for the 31 post-therapy samples, with a median of 149 (Supplementary Figure 1 (B)).

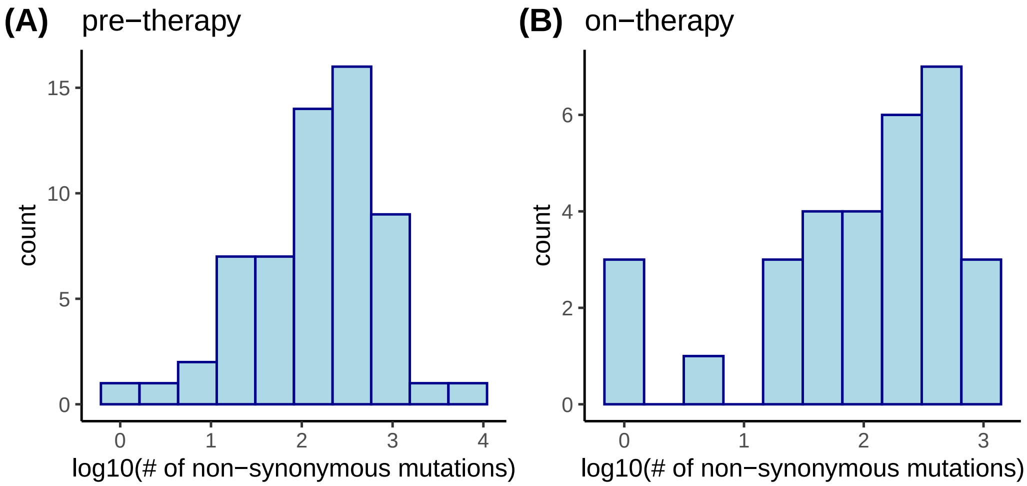

Supplementary Figure 2: The distribution of log10(# of non-synonymous mutations) for 59 pre-therapy samples and 31 on-therapy samples.

#### C.2 Impute HLA genotypes

We imputed HLA genotypes by OptiType (`https://github.com/FRED-2/OptiType`), an HLA genotyping algorithm based on integer linear programming (Szolek et al., 2014). OptiType took the FASTQ files as input, mapsped the reads to a customized HLA allele reference, and genotyped the HLA at four digit resolution. Compared to other *in silico* approaches, OptiType had significantly better performance. On the 361 benchmark samples, OptiType had an overall accuracy of 97.1%, outperforming other methods on all datasets by 4-15% accuracy. Consequently, OptiType had a 65-83% lower rate of incorrect allele predictions compared to other methods.

##### C.3 Prioritizing Neoantigens

First, for each non-synonymous SNV, we extracted all possible 9AA peptides that covered the specific SNV. We implemented the following procedure. Each non-synonymous SNV was annotated to a transcript with a unique ensembl transcript ID. Its location in the corresponding protein and the amino acid change (the `AAChange.ensGene` field of ANNOVAR output) was recorded in the format like 'p.G12C', meaning the 12-th amino acid is a Glutamine in the wild type protein and a Cysteine in the mutated protein. We extracted the protein sequence of each transcript by the R function `getSequence` in the `biomaRt` library and replaced the wild-type amino acid with the mutated amino acid to obtain mutated peptides.

Then, for each tumor sample, we assessed the association between each mutated 9 AA peptide and each HLA by PEPPRMINT or NetMHCpan-4.1. Each somatic mutation was covered by at most 9 peptides. The number of peptides was smaller if the mutation was close to the beginning or the end of the protein. Each sample had at most 6 HLA-I alleles, or more precisely, 3, 3, 14, and 70 samples have 3, 4, 5, and 6 unique HLA-I alleles, respectively. Therefore, each somatic mutation had up to $9 \times 6 = 54$ association scores. We took the maximum as the score for the somatic mutation.

##### C.4 Additional Results

For PEPPRMINT, we ran all 15 neural networks to calculate the maximum scores per somatic mutation and the results were highly consistent across the 15 models (Supplementary Figure 2). We took the average across the 15 neural networks.

We also compared the neoantigen burden of pre-therapy samples vs. on-therapy samples, using 0.9 as the cutoff for the neoantigen score (Supplementary Figure 3). The patients who responded to the immunotherapy tended to have much lower

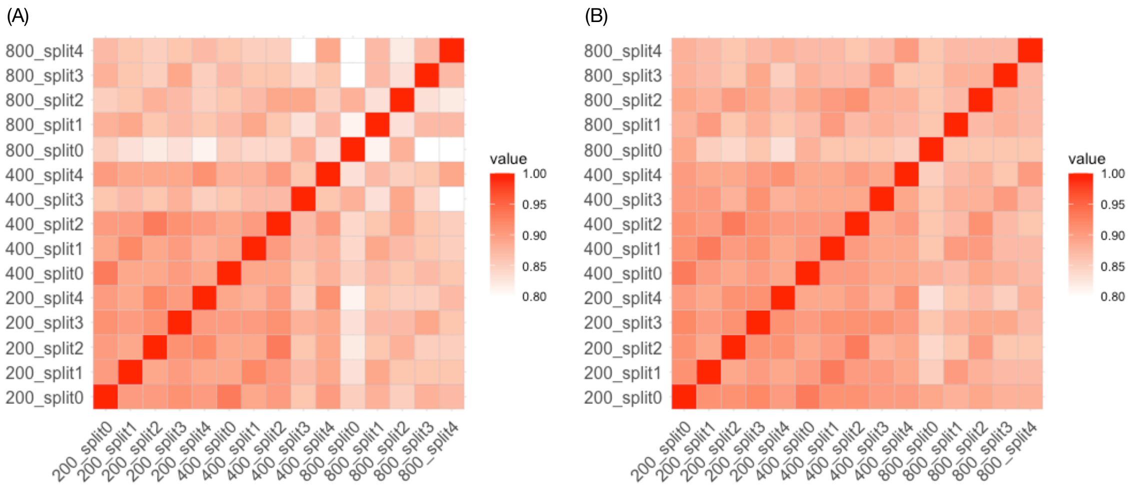

Supplementary Figure 3: Correlation of neoantigen scores for 31,298 somatic mutations across 15 neural networks using mutated peptides (A) or wild type peptides (B).

neoantigen burdens in the on-therapy sample than the pre-therapy samples. However, since only a subset of the patients (30 out of 60) had both pre-therapy and on-therapy samples, the sample size was too small for definite conclusions.

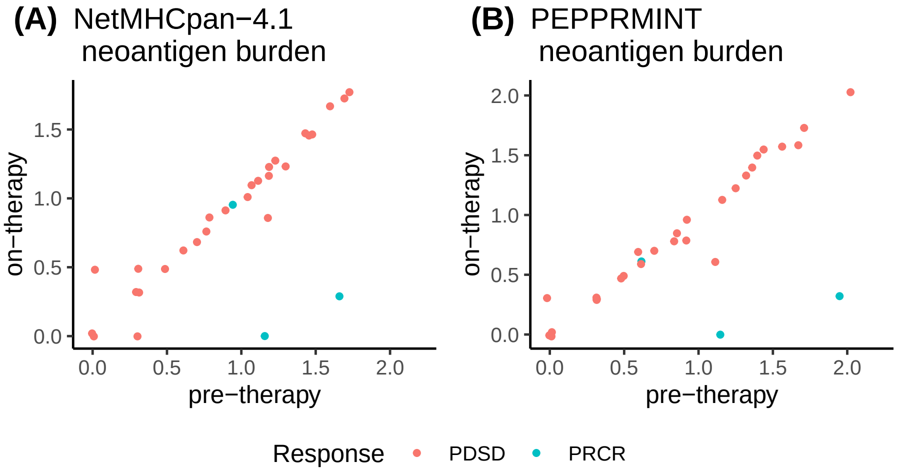

Supplementary Figure 4: Neoantigen burden comparison of pre-therapy samples vs. on-therapy samples. PDSD: progressive disease or stable disease. PRCR: partial response or complete response.

#### References

Alvarez, B., Reynisson, B., Barra, C., Buus, S., Ternette, N., Connelley, T., Andreatta, M., and Nielsen, M., 2019. NNAlign_MA; MHC peptidome deconvolution for accurate MHC binding motif characterization and improved T cell epitope predictions. *Molecular & Cellular Proteomics*, :in press.

Andreatta, M., Alvarez, B., and Nielsen, M., 2017. GibbsCluster: unsupervised clustering and alignment of peptide sequences. *Nucelic Acids Research*, **45**:W458–W463.

Andreatta, M., Lund, O., and Nielsen, M., 2013. Simultaneous alignment and clustering of peptide data using a gibbs sampling approach. *Bioinformatics*, **29**(1):8–14.

Bassani-Sternberg, M. and Gfeller, D., 2016. Unsupervised HLA peptidome deconvolution improves ligand prediction accuracy and predicts cooperative effects in peptide–HLA interactions. *The Journal of Immunology*, **197**(6):2492–2499.

Bulik-Sullivan, B., Busby, J., Palmer, C. D., Davis, M. J., Murphy, T., Clark, A., Busby, M., Duke, F., Yang, A., Young, L., *et al.*, 2019. Deep learning using tumor HLA peptide mass spectrometry datasets improves neoantigen identification. *Nature biotechnology*, **37**(1):55.

Cibulskis, K., Lawrence, M. S., Carter, S. L., Sivachenko, A., Jaffe, D., Sougnez, C., Gabriel, S., Meyerson, M., Lander, E. S., and Getz, G., *et al.*, 2013. Sensitive detection of somatic point mutations in impure and heterogeneous cancer samples. *Nature biotechnology*, **31**(3):213–219.

Henikoff, S. and Henikoff, J. G., 1992. Amino acid substitution matrices from protein blocks. *Proceedings of the National Academy of Sciences*, **89**(22):10915–10919.

Krøigård, A. B., Thomassen, M., Lænkholm, A.-V., Kruse, T. A., and Larsen, M. J., 2016. Evaluation of nine somatic variant callers for detection of somatic mutations in exome and targeted deep sequencing data. *PloS one*, **11**(3):e0151664.

Lek, M., Karczewski, K. J., Minikel, E. V., Samocha, K. E., Banks, E., Fennell, T., O'Donnell-Luria, A. H., Ware, J. S., Hill, A. J., Cummings, B. B., *et al.*, 2016. Analysis of protein-coding genetic variation in 60,706 humans. *Nature*, **536**(7616):285–291.

Li, H. and Durbin, R., 2009. Fast and accurate short read alignment with burrows–wheeler transform. *bioinformatics*, **25**(14):1754–1760.

Nielsen, M. and Andreatta, M., 2016. Netmhcpan-3.0; improved prediction of binding to mhc class i molecules integrating information from multiple receptor and peptide length datasets. *Genome medicine*, **8**(1):1–9.

Nielsen, M. and Andreatta, M., 2017. Nnalign: a platform to construct and evaluate artificial neural network models of receptor–ligand interactions. *Nucleic acids research*, **45**(W1):W344–W349.

Nielsen, M., Lundegaard, C., Blicher, T., Lamberth, K., Harndahl, M., Justesen, S., Røder, G., Peters, B., Sette, A., Lund, O., *et al.*, 2007. NetMHCpan, a method for quantitative predictions of peptide binding to any HLA-A and-B locus protein of known sequence. *PloS one*, **2**(8):e796.

O'Donnell, T. J., Rubinsteyn, A., Bonsack, M., Riemer, A. B., Laserson, U., and Hammerbacher, J., 2018. MHCflurry: open-source class I MHC binding affinity prediction. *Cell systems*, **7**(1):129–132.

O'Donnell, T. J., Rubinsteyn, A., and Laserson, U., 2020. MHCflurry 2.0: Improved pan-allele prediction of MHC class I-presented peptides by incorporating antigen processing. *Cell systems*, **11**(1):42–48.

Racle, J., Michaux, J., Rockinger, G., Arnaud, M., Bobisse, S., Chong, C., Guillaume, P., Coukos, G., Harari, A., Jandus, C., *et al.*, 2019. Robust prediction of HLA class II epitopes by deep motif deconvolution of immunopeptidomes. *Nature Biotechnology*, :in press.

Reynisson, B., Alvarez, B., Paul, S., Peters, B., and Nielsen, M., 2020. Netmhcpan-4.1 and netmhciipan-4.0: improved predictions of mhc antigen presentation by concurrent motif deconvolution and integration of ms mhc eluted ligand data. *Nucleic Acids Research*, .

Saunders, C. T., Wong, W. S., Swamy, S., Becq, J., Murray, L. J., and Cheetham, R. K., 2012. Strelka: accurate somatic small-variant calling from sequenced tumor–normal sample pairs. *Bioinformatics*, **28**(14):1811–1817.

Szolek, A., Schubert, B., Mohr, C., Sturm, M., Feldhahn, M., and Kohlbacher, O., 2014. Optitype: precision hla typing from next-generation sequencing data. *Bioinformatics*, **30**(23):3310–3316.

Van der Auwera, G. A. and O'Connor, B. D., 2020. *Genomics in the Cloud: Using Docker, GATK, and WDL in Terra*. O'Reilly Media.

Wang, K., Li, M., and Hakonarson, H., 2010. ANNOVAR: Functional annotation of genetic variants from next-generation sequencing data . *Nucleic Acids Research*, **38**:e164.

Xu, H., DiCarlo, J., Satya, R. V., Peng, Q., and Wang, Y., 2014. Comparison of somatic mutation calling methods in amplicon and whole exome sequence data. *BMC genomics*, **15**(1):1–10.

Zhao, W. and Sher, X., 2018. Systematically benchmarking peptide-mhc binding predictors: From synthetic to naturally processed epitopes. *PLoS computational biology*, **14**(11):e1006457.
